# *TTG2* defines trichome cell shape by modulating microtubules and actin cytoskeleton in Arabidopsis

**DOI:** 10.1101/2023.03.29.534767

**Authors:** Lu Liu, Weihua Cao, Lanxin Yuan, Lan Yang, Yali Wang, Chi Zhang, Dan Wang, Wenjia Wang, Hongchang Zhang, John Schiefelbein, Fei Yu, Lijun An

## Abstract

The Arabidopsis *TRANSPARENT TESTA GLABRA2* (*TTG2*) gene encodes a WRKY transcription factor that regulates a range of development events like trichome, seed coat, and atrichoblasts formation. Loss-of-function of *TTG2* was previously shown to reduce or eliminate trichome branching. Here, we report the identification of a new allele of *TTG2*, *ttg2-6*. In contrast to the *ttg2* mutants described before, *ttg2-6* displays unique trichome phenotypes. Some *ttg2-6* mutant trichomes are hyperbranched, whereas others are hypobranched, distorted, or clustered. Further, we found that in addition to specifically activating R3 MYB transcription factor *TRIPTYCHON* (*TRY*) to promote trichome branching, *TTG2* also integrates cytoskeletal signaling to regulate trichome morphogenesis. The *ttg2-6* trichomes display aberrant cortical microtubules (cMTs) and cortical actin filaments (F-actin) configurations. Moreover, genetic and biochemical analysis showed that TTG2 could directly bind and regulate the expression of *BRICK1* (*BRK1*), a subunit of the actin nucleation promoting complex SCAR/WAVE. Collectively, taking advantage of *ttg2-6*, we uncovered new functions for *TTG2* in facilitating cMT and F-actin cytoskeleton-dependent trichome development. Our evidence for a direct relationship between *TTG2* and cytoskeletal regulators establishes an unprecedented understanding of cellular signaling events downstream of the core transcriptional regulation during trichome development in Arabidopsis.

## Introduction

Trichomes are specialized epidermal structures in plants, and have long been favored as a model system to elaborate cell differentiation and morphogenesis in plants due to their unique developmental patterns (Hülskamp et al., 2004). In Arabidopsis, the leaf trichomes are single-celled structures, and usually exhibit a well-extended stalk crowned with three or four straight branches (Hülskamp et al., 1994; Folkers et al., 1997). They are also well arranged in spatial distributions, and clustered trichomes are rare (Hülskamp et al., 1994). The formation of such unique patterning forms and architecture depend on a group of key genetic factors.

*TRANSPARENT TESTA GLABRA 2* (*TTG2*) encodes a WRKY transcription factor that plays important roles during trichome development (Johnson et al., 2002). Inactivation of *TTG2* leads to pleiotropic trichome developmental defects including clustered trichomes, reduced trichome density, and abnormal trichome branches (Johnson et al., 2002), implying *TTG2* may participate in multiple regulatory pathways. The temporal and spatial expression of *TTG2* is directly regulated by the transcriptional activation complex comprised of the R2R3 MYB protein GLABRA 1 (GL1), the basic helix-loop-helix (bHLH) proteins GLABRA 3 (GL3) and ENHANCER OF GL3 (EGL3), and the WD repeat protein TRANSPARENT TESTA GLABRA 1 (TTG1) (Johnson et al., 2002; Ishida et al., 2007; Zhao et al., 2008). All of these are considered to be the first tier positive regulators for trichome patterning (Morohashi and Grotewold, 2009), and mutation of each of them results in glabrous or decreased trichome numbers (Ishida et al.,2008; Tominaga-Wada et al., 2011; Han et al., 2022). However, the downstream pathways of TTG2 during trichome development are unclear so far. The only direct target of *TTG2* identified is the patterning gene, *TRIPTYCHON* (*TRY*), which encodes a R3 MYB transcription factor (Hülskamp et al., 1994; Schellmann et al., 2002; Pesch et al., 2014). TTG2 physically binds to the promoter of *TRY* to active *TRY* expression (Pesch et al., 2014). The *try* mutant shows a cluster frequency and reduced trichome numbers that is similar to that in the *ttg2-1* mutant (Hülskamp et al.,1994; Johnson et al., 2002), implicating *TTG2* as a regulator of *TRY* for trichome initiation and distribution. Given the multiple trichome defects in *ttg2* mutants, it is interesting to consider the possibility that other cellular pathways coordinate with *TTG2* to modulate trichome development.

Genetic and pharmacological information have implicated the distinct requirements of cortical microtubules (cMTs) and cortical actin filaments (F-actin) cytoskeleton organization in defining trichome development (Ishida et al., 2008; Tominaga-wada et al., 2011; Li et al., 2019). Specifically, it has been suggested that microtubules establish the formation and spatial distribution of trichome branches whilst actin microfilaments elaborate and maintain the overall trichome branch shapes (Mathur et al., 1999; Sambade et al., 2014). During trichome branching, microtubules reorient from a transverse to a longitudinal direction with respect to the branch growth axis (Mathur and Chua, 2000). Meanwhile, actin microfilaments assume an increasing degree of complexity from fine filaments to thick, longitudinally stretch cables during trichome formation (Mathur et al., 1999; Schwab et al., 2003). Consistently, mutations in genes involved in cMTs and F-actin assembling and dynamics often lead to aberrant branch numbers and shapes (Tominaga-Wada et al., 2011; Yanagisawa et al., 2015; Liang et al., 2019). *ZWICHEL* (*ZWI*) encodes a Kinesin-like Calmodulin Binding Protein (KCBP) (Oppenheimer et al., 1997). Mutation in the *ZWI* results in trichomes that failed to branch properly, causing less-branched trichomes with shortened stalks and blunted branch ends (Hülskamp et al.,1994; Oppenheimer et al., 1997). Further studies demonstrated that ZWI may serve as a hub to integrate and coordinate cMTs and F-actin to achieve the cytoskeletal configuration necessary for trichome development (Tian et al., 2015).

The *DISTORTED* group genes encode subunits of the actin-related protein (ARP)2/3 and suppressor of cyclic AMP repressor (SCAR)/Wiskott-Aldrich syndrome protein family verprolin homologous protein (WAVE) complexes (Ishida et al., 2008; Tominaga-Wada et al., 2011; Li et al., 2019), and loss-of-function of mutants possess fascinating “distorted” trichomes with alterations in the organization and/or density of F-actin in expanding trichomes (Hülskamp et al.,1994; Schwab et al., 2003). HSPC300 is a conserved 8 kDa human protein and participates in ARP2/3 activation through interacting with WAVE proteins (Stovold et al., 2005). *BRICK1* (*BRK1*) is the maize homolog of *HSPC300* and loss-of-function mutations result in the absence of epidermal pavement cell lobes formation and short and swollen leaf hairs (Frank and Smith, 2002; Gallagher and Smith, 2000). These defects were associated with loss of localized cortical F-actin enrichment, implying a role for *BRK1* in promoting actin polymerization (Frank and Smith, 2002; Gallagher and Smith, 2000). Likewise, the Arabidopsis *brk1* mutant displays a strongly distorted trichome phenotype accompanied by the alteration in F-actin organization in trichome branches (Djakovic et al., 2006). *BRK1* is also reported essential for the SCAR/WAVE-ARP2/3 pathway in plants by selectively stabilizing SCAR1 and SCAR2 proteins (Djakovic et al., 2006; Le et al., 2006). However, the signals responsible for regulating *BRK1* activity are unknown.

In this study, we report that the WRKY transcription factor TTG2 directly binds and regulates the expression of *BRK1* during trichome morphogenesis. We found that a new mutant allele of *TTG2*, called *ttg2-6*, produces conspicuous distorted trichomes which are reminiscent of *brk1* mutants. Genetic and molecular analyses indicated that *BRK1* functions downstream of *TTG2*. Moreover, we showed that TTG2 directly binds to the W-box in the *BRK1* promoter region. Taken together, our results suggest that *TTG2* facilitates cytoskeleton-dependent trichome development, uncovering a new pathway in trichome development as well as in *BRK1* regulation.

## Results

### *abt4-1* mutant displays pleiotropic trichome developmental defects

In our continuous effort to uncover novel molecular mechanisms involved in plant cell patterning, initiation, and morphogenesis, we performed genetic screening for trichome phenotypes, and a unique recessive trichome mutant, designated as *<u>a</u>berrantly <u>b</u>ranched trichome 4-1* (*abt4-1*) was obtained. The overall growth and development of *abt4-1* was not significantly different from that of wild-type (WT) except for larger cotyledons, but it exhibited striking pleiotropic trichome defects (Figure 1, A-G; Supplemental Figure S1). Under our growth conditions, the trichomes of WT plants are predominately three- or four-branched (approximate 71% three-branched and 28% four-branched on the 3^rd^ rosette leaves, respectively), while less or more branched trichomes are rare (Figure 1, A, C, D, and H).

**Figure 1.**
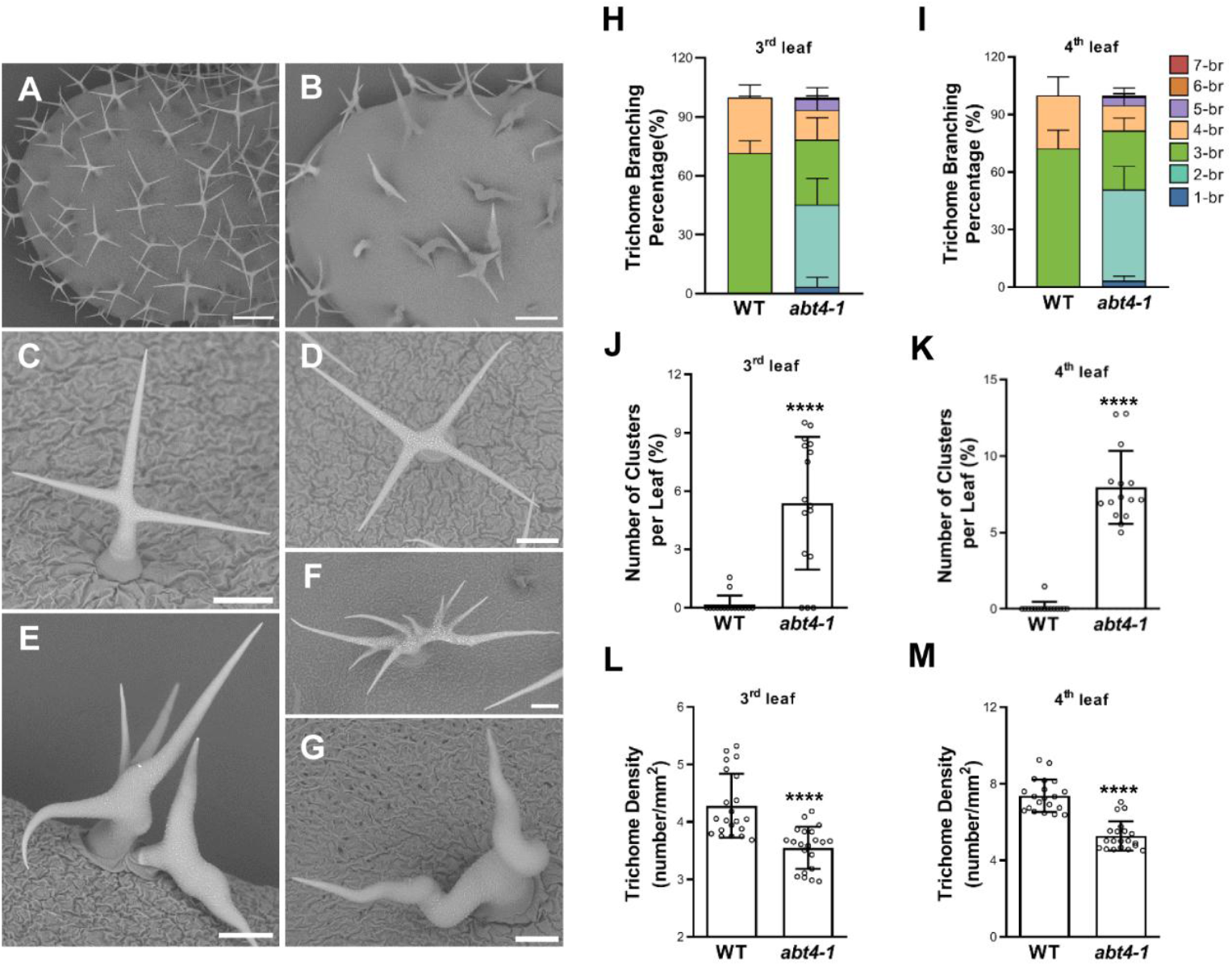
Characterization of trichome phenotypes from the wild type (WT) and *abt4-1* mutant plants. A-G, Representative trichomes on the fifth rosette leaves of three-week-old WT (A, C and D) and *abt4-1* (B, and E-G). For A and B, bars=400 μm, for C-G, bars=100 μm. H-M, Quantification of the trichome branch numbers (H and I), cluster numbers (J and K), and trichome density (L and M) of in WT and *abt4-1* on the third (H, J and L) and fourth rosette leaves (I, K and M) respectively. 1/2/3/4/5/6/7-br for one/two/three/four/five/six/seven-branched trichomes, respectively. Data are shown as means ± SD. ****P ≤ 0.0001 by the Student’s t test.

Moreover, WT trichomes are precisely spaced on leaves and clustered trichomes are scarcely observed (Figure 1, A and J). In contrast, *abt4-1* mutants displayed trichome developmental defects in multiple aspects. First, *abt4-1* trichomes were variable in branch numbers, ranging from one- to seven-branched, and even nine-branched trichomes were observed (Figure 1, E-G, and H). Second, some of the trichomes in *abt4-1* showed twisted and swollen branches (Figure 1, E and G), resembling to those of dis-group mutants (Hülskamp et al., 1994; Schwab et al., 2003). Third, *abt4-1* mutants also exhibited trichome distribution defects, with ∼5% of trichomes arising closely to another and forming trichome clusters, as compared to only 0.1% in WT (Figure 1, E and J). Fourth, trichome density was apparently decreased in *abt4-1* plants (approximately 2.9 trichomes per mm^2^ in *abt4-1* versus 3.6 trichomes per mm^2^ in WT) (Figure 1L), implying that trichome initiation was also impaired. Finally, some *abt4-1* trichomes lacked surface papillae, and appeared dark gray when viewed under scanning electron microscopy (Figure 1B), indicating a defective trichome maturation process. Similar trichome developmental differences were also observed in the 4^th^ rosette leaves between the WT and the *abt4-1* mutant (Figure 1, I, K, and M). Taken together, these phenotypic observations suggest that *ABT4* serves as a crucial genetic factor controlling multiple aspects of trichome development.

### *abt4-1* is a new mutant allele of *TTG2*

Trough map-base cloning and nucleotide sequencing, we found a G to A substitution in the 4^th^ exon of *AT2G37260* gene in *abt4-1* (Figure 2A). *AT2G37260* encodes the key regulator of trichome development, TTG2 (Johnson et al., 2002). Theoretically, this mutation would convert the corresponding codon from cysteine to tyrosine (C379Y) that is located in the second CCHH zinc finger motif in TTG2 (Figure 2B), which is assumed to take part in target binding and is well conserved in *TTG2* homologs (Johnson et al., 2002; Li et al., 2015). These results indicated that the *abt4-1* trichome defects may result from the mutation of *TTG2*.

**Figure 2.**
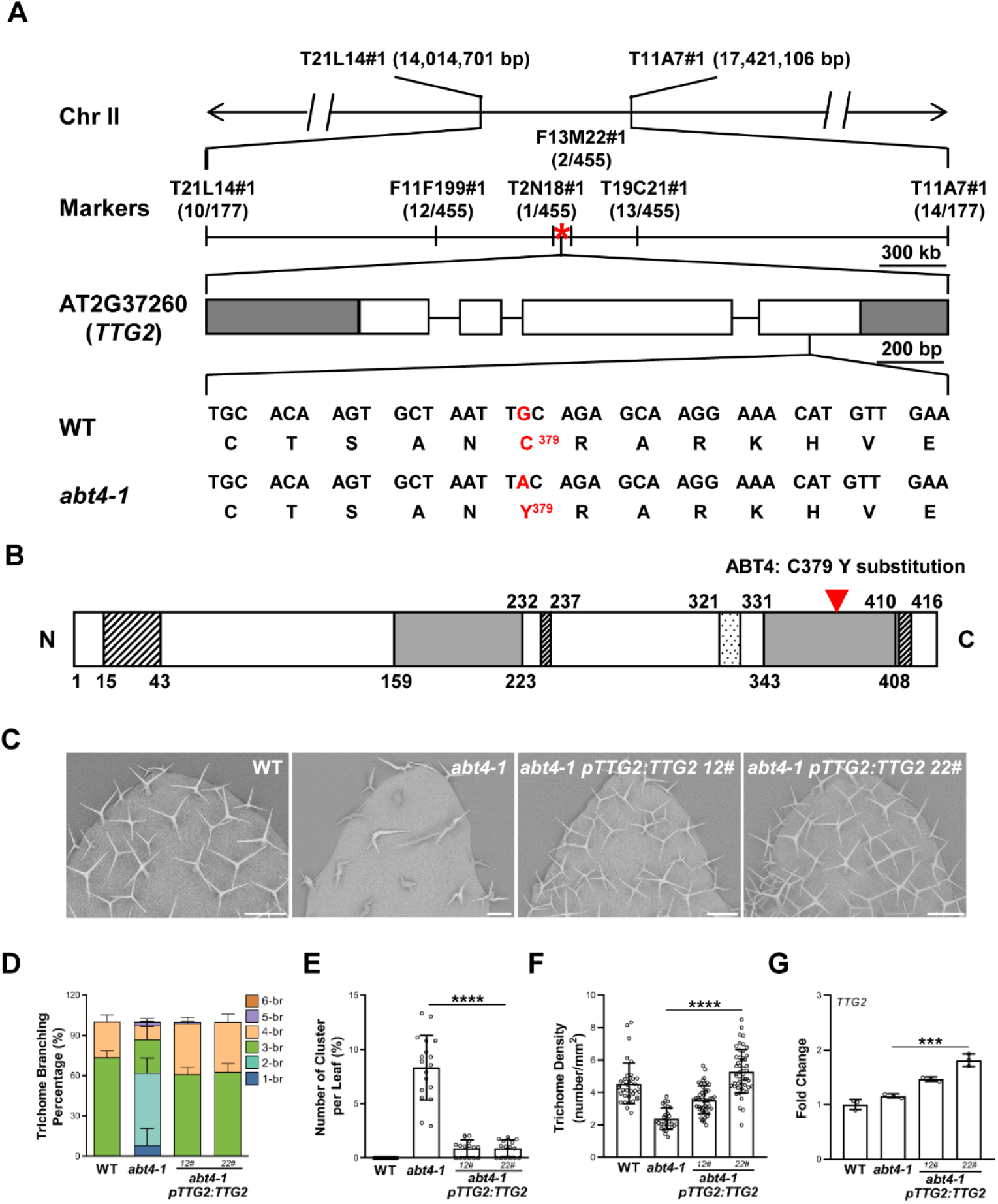
Cloning and genetic complementation of *ABT4*. A, Map-based cloning of *abt4-1*. The *ABT4* locus was preliminarily located between markers T21L14 and T11A7 on chromosome II. Fine-mapping restricted the locus within a region between T2N18 and F13M22. The numbers of genetic recombinants are shown under each marker. The asterisk indicates the position of the *TTG2* gene, *AT2G37260*. In the schematic gene structure, white rectangles and solid lines represent the exons and introns, respectively. The 5’and 3’untranslated regions are shown as gray rectangles. The highlighted nucleotide below the gene model indicates the exact position of the mutation site. B, TTG2 protein structure and deduced mutation in the amino acid sequence. The two WRKY domains are indicated as shaded boxes. The putative Ser/Thr-rich activation domains and a possible nuclear localization signal are shown as slash lined and dotted boxes, respectively. The red arrowhead shows the position of the C379Y mutation in ABT4 located in the second WRKY domain. C, Trichome branching phenotypes of WT, *abt4-1* and the genetic complementation lines (*abt4-1 pTTG2:TTG2 12#* and *abt4-1 pTTG2:TTG2 22#*). Bars=100 μm. D-F, Quantification of trichome branch numbers (D), cluster number (E) and trichome density (F) on the fourth rosette leaves of plants shown in C. 1/2/3/4/5/6-br for one/two/three/four/five/six-branched trichomes, respectively. Data are presented as mean ±SD. ****P ≤ 0.0001by the Student’s t test. G, RT-qPCR analysis of *TTG2* transcripts levels in WT, *abt4-1* and the complementation lines. Data are shown as means ±SD of three biological replicates. ****P ≤ 0.0001 and ***P ≤ 0.001 by the Student’s t test.

Next, we performed genetic complementation experiments to determine whether *ABT4* is represented by *TTG2*. After expressing the coding region of *TTG2* under the control of its native promoter (*pTTG2:TTG2*) in the *abt4-1* genetic background, we identified independent *abt4-1 pTTG2:TTG2* transgenic lines that showed higher expression levels compared to that in WT (Figure 2G). Consistently, these plants produced WT-like trichome phenotypes including morphology, distribution, and trichome density (Figure 2F). Quantitative measurements confirmed the phenotypic observations (Figure 2, C-E), indicating that *in situ* expression of *TTG2* can rescue the trichome developmental defects of *abt4-1*.

In previous reports describing the *ttg2* mutants in the L*er* background (*ttg2-1* and *ttg2-2*), a major defect is to reduce or eliminate branching (Johnson et al., 2002), which is in strikingly contrast to the hyper-branched trichomes observed in *abt4-1*. To examine this issue, we identified two independent T-DNA insertion lines of *TTG2* in the Col background, *Salk_148838* (*ttg2-3*) and *Salk_206852* (*ttg2-7*) (Figure 3A). These two mutations likely represent null alleles of *TTG2* because we did not detect any *TTG2* transcripts in both *ttg2-3* and *ttg2-7* (Figure 3F). Phenotypically, homozygous *ttg2-3* and *ttg2-7* mutants showed similar defects as those in *abt4-1*, including reduced trichome numbers, and clustered and distorted trichomes (Figure 3, B, D, and E). However, unlike the variable trichome branch numbers in *ttg2-6*, the majority of trichomes were one- or two-branched in *ttg2-3* and *ttg2-7* (Figure 3, B and C). The F1 plants of crosses between *ttg2-6* and *ttg2-3,* as well as *ttg2-6* and *ttg2-7* were defective in trichome numbers, distributions, and branch extensions (Figure 3, B, D, and E), indicating that both *ttg2-3* and *ttg2-7* failed to complement *ttg2-6* for these defects.

**Figure 3.**
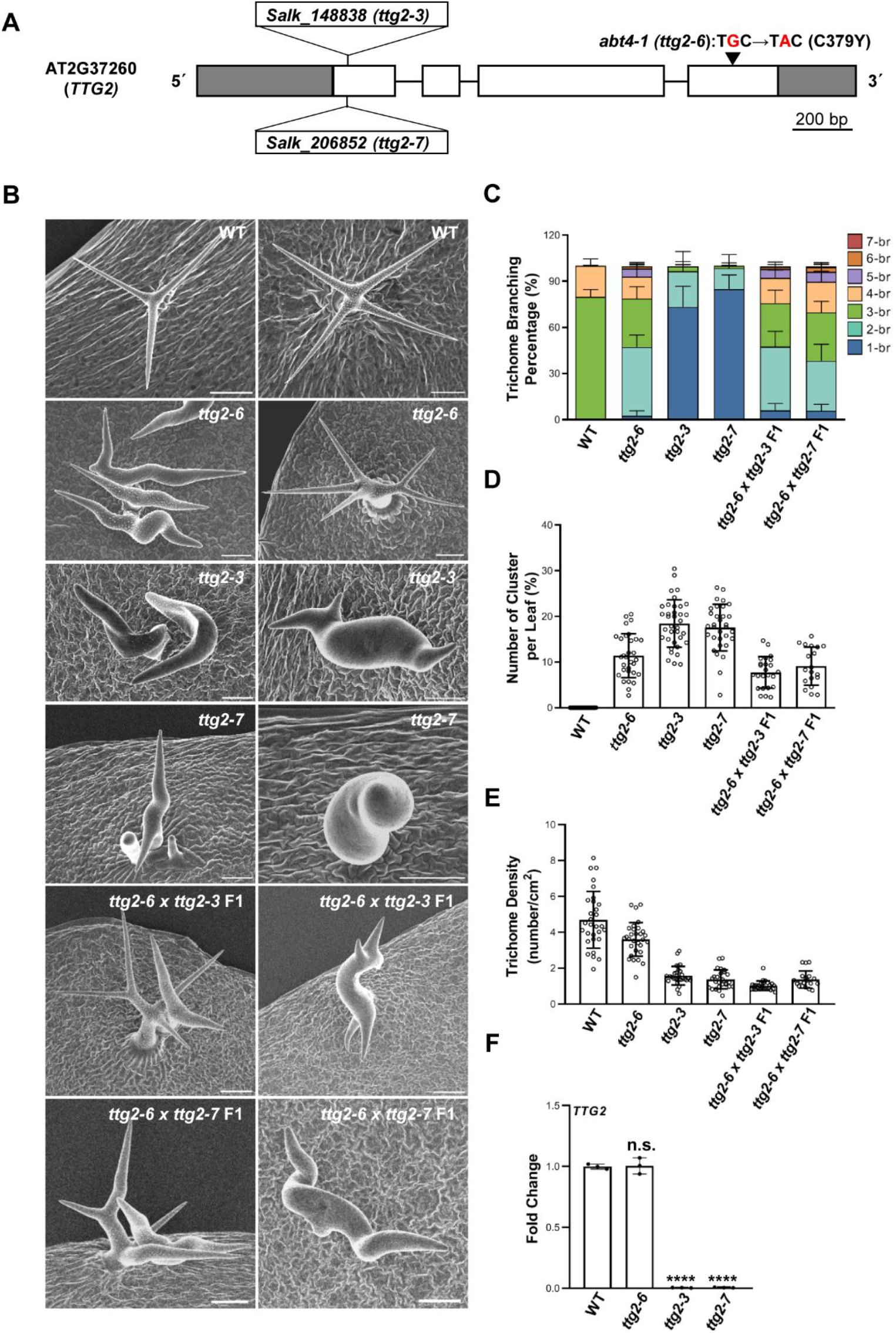
Trichome phenotypes of various *TTG2* alleles. A, Schematic representation of the mutation sites in *abt4-1, ttg2-3* (*Salk_148838*), and *ttg2-7* (*Salk_206852*). B, Representative trichomes on the fifth rosette leaves of WT, *abt4-1*, *ttg2-3*, and *ttg2-7*. Bars=100 μm. C-E, Quantification of trichome branch numbers (C), cluster numbers (D), and trichome density (E) of WT, *abt4-1*, *ttg2-3*, *ttg2-7*, *abt4-1x ttg2-3* F1, and *abt4-1x ttg2-7* F1. Data were shown as means ±SD. **P ≤ 0.01 by the Student’s t test. F, The relative expression levels of *TTG2* in different *TTG2* alleles measured by RT-qPCR. Data are shown as means ± SD of three biological replicates. n.s. no significance, ****P ≤ 0.0001 by the Student’s t test.

Nevertheless, the trichome branching pattern of the F1 plants resembled that of *ttg2-6* (Figure 3, B and C), suggesting *ttg2-6* is epistatic to *ttg2-3* and *ttg2-7* with regard to the degree of trichome branching. Among the mutant alleles of *TTG2* reported so far (Johnson et al., 2002; Ishida et al., 2007), only *ttg2-6* displayed hyper-branched trichomes. Thus, *ttg2-6* is likely to represent a unique allele of *TTG2*. Taken together, these results demonstrate that *ABT4* is *TTG2*, and the mutation in *TTG2* results in the trichome defects of *abt4-1*. Therefore, *abt4-1* was renamed as *ttg2-6* based on earlier nomenclature.

### Altered cytoskeletal configuration in ***ttg2-6*** mutant

Given the highly similar trichome morphologies of the *ttg2-6* mutant and those of loss-of-function mutants of genes encoding microtubule- and actin-associated proteins (Ishida et al., 2008; Tominaga-wada et al., 2011; Li et al., 2019), we reasoned that *TTG2* might regulate trichome development through the cytoskeletons. To address this issue, we first examined the cMTs and F-actin configurations in mature trichomes. As shown in Figure 4, the cMTs arrays in WT trichomes were assembled in a highly ordered fashion and laid out along with the extension directions of stalks and branches as previously reported (Mathur and Chua, 2000; Figure 4A). However, in *ttg2-6* background, the cMTs were relatively thin and diffused and numerous microtubules crisscrossed in both stalks and branches (Figure 4A). Notably, a portion of transverse cMTs were retained in stalks (Figure 4A). These results thus established a certain degree of correlation between the trichome morphologies of *ttg2-6* and the cMT arrangements. To further elaborate the influence of *TTG2* mutation on the cMTs arrangements, we quantitatively analyzed their configurations. Consistent with our visual observations, the angles of cMTs were arranged from 0° to 15° in WT (Figure 4B), indicating an approximately parallel orientation to the growth direction. However, in the *ttg2-6* background, the angles of cMTs displayed a more scattered distribution from 15° to 90° with a significant peak from 75° to 90° (Figure 4B). We also examined the anisotropy of the cMTs in *ttg2-6*, and found that the mean of the anisotropy value in *ttg2-6* was about 0.18±0.08, compared with that of about 0.47±0.07 in WT (Figure 4C), suggesting an isotropic arrangement of cMTs in *ttg2-6* comparing to WT. The organization of the cMT arrays in the pavement cells in *ttg2-6* was also examined. Strikingly, the organization of the cMTs in *ttg2-6* was indistinguishable from that of WT (Supplemental Figure S3, A and B), implying that the impact of *TTG2* on cMTs dynamics was probably trichome specific.

**Figure 4.**
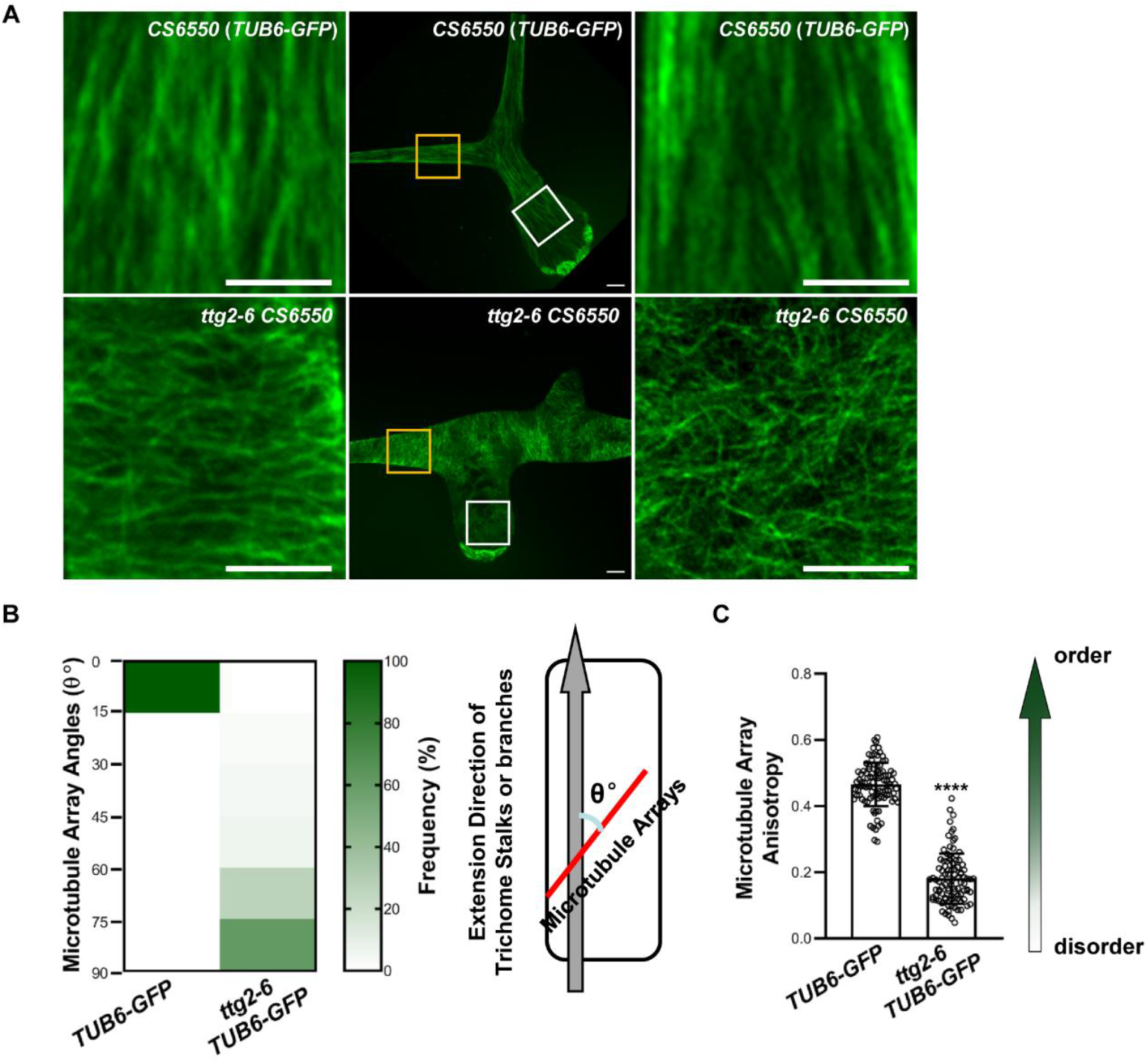
Cortical microtubule (cMT) organization in *ttg2-6* mature trichomes. A, Representative microtubule arrays in the branches (left panel) and stalks (right panel) of mature trichome in WT and *ttg2-6* mutant observed under spinning disk confocal microscopy. The yellow and white bordered boxes indicate the trichome branches and stalks, respectively. The images in the left and right panel are enlarged version of the boxed parts in the middle photos. Bars=20 μm. B and C, Quantitative analysis of the cMT phenotypes with respect to cMT angles (B) and anisotropy measurements (C). The angle of microtubules that are parallel to the stalk and branch extension directions is defined as 0°, whereas the angle of those perpendicular to the growth axis is defined as 90°. Approximately 1000 microtubules angles were measured for each genotype, and then these were divided into six intervals (0°-15°, 15°-30°, 30°-45°, 45°-60°, 60°-75°, 75°-90°). The frequency in each interval was calculated by dividing the number of microtubules with the corresponding angles in each interval by the total number of microtubules, and then displaying as a heatmap with GraphPad Prism software. For anisotropy evaluation, about 50 independent trichomes were used for analysis, and two independent areas for each trichome were selected for anisotropy measurements. According to Boudaoud et al. (2014), the anisotropy score zero is defined for purely isotropic arrays (no order) and one is defined for purely anisotropic arrays (perfectly order). Data are shown as means ± SD. ****P ≤ 0.0001 by the Student’s t test.

The F-actin organization in mature trichomes was also disrupted in *ttg2-6* mutant. As reported previously, we observed that F-actin arrays in WT trichomes were fairly well aligned with each other along the long axes of trichome branches, and they were relatively uniformly distributed (Figure 5A). Quantitative analysis showed that angles of actin filaments were arranged from 0° to 30°, and the mean value of the anisotropy was 0.44±0.1 (Figure 5, B and C). In *ttg2-6* mutants, the situation was more complex. In those trichomes which were overbranched but with normal branch morphology, the organizations of the F-actin arrays were indistinguished from that in WT in both stalks and branches. While in those distorted trichomes, the organization of F-actin arrays was profoundly altered, such that filaments were almost aligned transversely to the axis of growth directions, and they were unevenly distributed with in the cells with areas of bright actin restricted to indentation regions of the trichomes (Figure 5A). Quantitative analysis showed that angles of the actin filaments were arranged from 30° to 90°, and the mean value of the anisotropy was 0.25±0.11 (Figure 5, B and C). These findings indicated a requirement of *TTG2* on F-actin configuration during trichome development and the alteration of F-actin arrays alignment and distributions might be associated with the swelling and distortion of mutant trichomes in *ttg2-6*.

**Figure 5.**
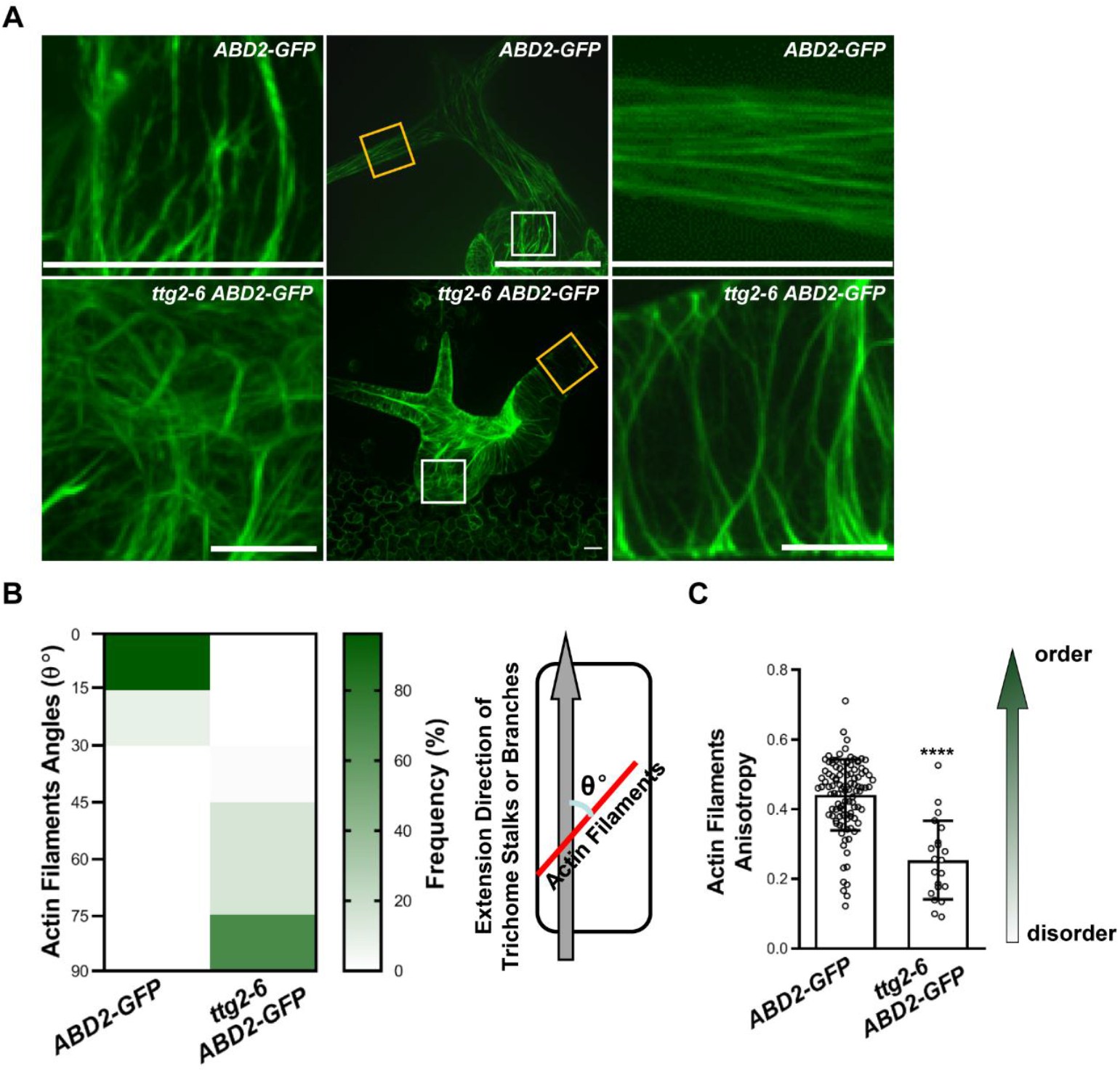
Cortical actin filaments (F-actin) arrangements in *ttg2-6* mature trichomes. A, Typical F-actin arrays in the stalks (left panel) and branches (right panel) of mature trichomes in WT and *ttg2-6* mutant observed under spinning disk confocal microscopy. The yellow and white bordered boxes indicate the trichome branches and stalks, respectively. The images in the left and right panel are enlarged version of the boxed parts in the middle pictures. Bars=20 μm. B and C, Quantification of F-actin arrangements with respect to F-actin angles (B) and anisotropy (C). Similar parameters and methods are employed as those used for the cMTs organization analysis. Data were shown as means ± SD. ****P ≤ 0.0001 by the Student’s t test.

To investigate whether the effects of *TTG2* on F-actin configuration is trichome specific, we monitored F-actin organization in expanding cotyledon pavement cells as well. Although the overall shape of the *ttg2-6* pavement cells was similar to that of WT, the actin configuration was different. The actin arrays were assembled more orderly compared to WT (Supplemental Figure S3C), with an average anisotropy value of 0.26±0.13 in *ttg2-6* versing 0.17±0.11 in WT (Supplemental Figure S3D) . In addition, some thick F-actin cables was also found in *ttg2-6* (Supplemental Figure S3C). These results indicated that the effects of *TTG2* on actin cytoskeleton might be globally.

### ***TTG2*** interacts genetically with ***BRK1*** and ***TTG2*** promotes ***BRK1*** expression

The striking effects of *TTG2* on cMTs and F-actin during trichome development raised the possibility that *TTG2* may influence cytoskeleton organization through targeting downstream genes regulating cMTs and F-actin. Therefore, we constructed a series of double mutants by crossing *ttg2-6* with mutants of genes encoding proteins involved in cMTs dynamics and SCAR/WAVE-ARP2/3 subunits, respectively (Supplemental Figures S4-S7). Since *ZWI* was reported to integrate both microtubule and actin cytoskeleton to determine trichome cell shape (Tian et al., 2015), we first examined the trichome phenotypes of *ttg2-6 zwi-101* double mutants. As shown in Supplemental Figure S4, *ttg2-6 zwi-101* displayed clustered, twisted trichomes with blunt ends, suggesting an additive interaction between *TTG2* and *ZWI*. During of investigating of trichome phenotypes of other double mutants, we found that *TTG2* interacts genetically with *BRK1*. *BRK1* mutations result in trichome branches that fail to elongate normally after initiation (Frank and Smith, 2002; Djakovic et al., 2006). In *ttg2-6 brk1- 1* double mutants, both the overall plant phenotypes and the trichomes branch morphologies were very similar to the *brk1-1* single mutant (Figure 6, A and B), indicating *brk1* was probably epistatic to *ttg2* in regulating branch extension. Consistently, *ttg2-6 brk1-1* trichomes also displayed similar actin cytoskeleton organization impairments to those of *brk1-1* (Figure 7, A and B). The parallel-aligned actin cables observed in WT trichomes were lost, and instead, a meshwork of transverse, intertwined actin filaments dominated in the *ttg2-6 brk1-1* trichomes (Figure 7A). Coincidently, as in *brk1-1*, quantitative analyses showed the F-actin angles were scattered from 15° to 90° in *brk1-1 ttg2-6* (Figure 7B), suggesting variable directions of the actin filaments. Meanwhile, we noticed that the genetic relationship between *BRK1* and *TTG2* was complex with respect to other trichome phenotypes. The branch numbers of *ttg2-6 brk1-1* trichomes were further decreased compared with the respective single mutants (Figure 6C), while the trichome density and distribution were reminiscent of that of *ttg2-6* (Figure 6, D and E). These results suggest that *TTG2* may interact with *BRK1* to specifically regulate trichome branch initiation and extension. In contrast, *TTG2* may integrate other pathways to control trichome differentiation and distribution.

**Figure 6.**
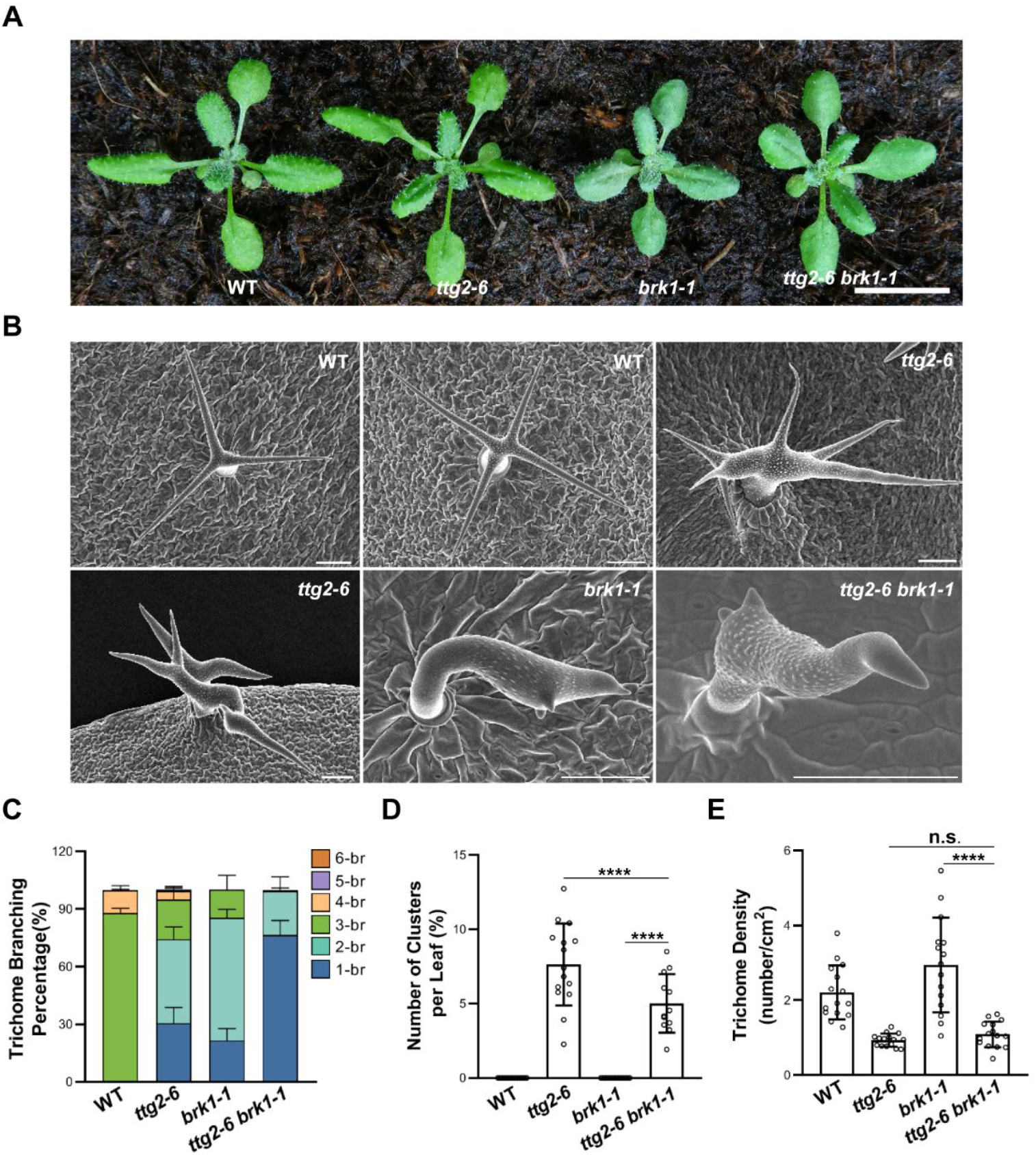
Phenotypes of *ttg2-6 brk1-1* double mutants. A, Overall plant morphology of two-week-old soil grown WT, *ttg2-6*, *brk1-1*, and *ttg2-6 brk1-1*, bar=1 cm. B, Representative trichomes on the fifth rosette leaves of WT, *ttg2-6*, *brk1-1*, and *ttg2-6 brk1-1*, bars=100 μm. C-E, Quantification of trichome branch numbers (C), cluster number (D), and trichome density (E) of the plants shown in panel B. Data are shown as means ±SD. n.s. no significance, ****P ≤ 0.0001 by the Student’s t test.

**Figure 7.**
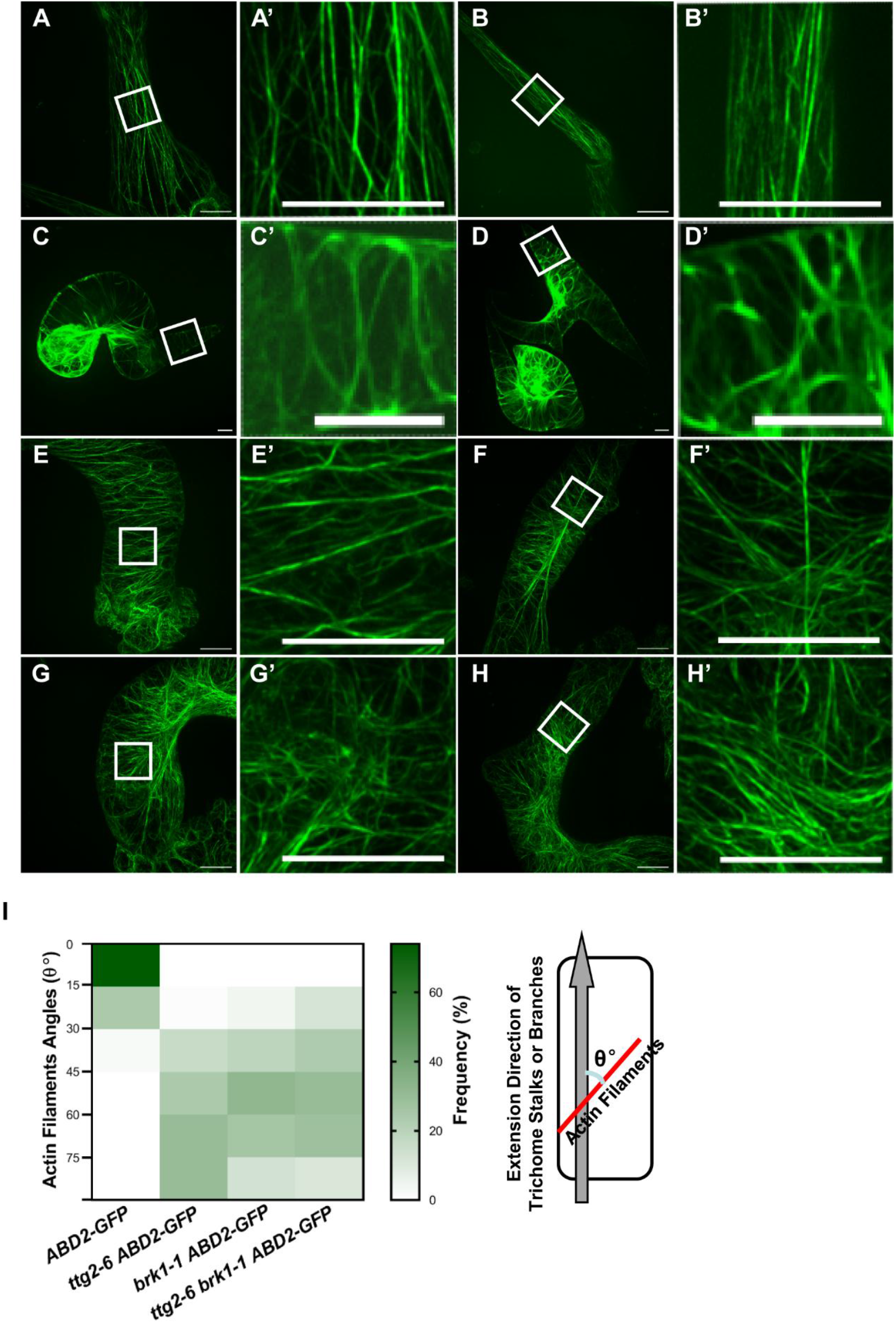
F-actin organization of *ttg2-6 brk1-1* double mutants. A, C, E, and G, Typical actin arrays in the stalk of mature trichomes in WT (A), *ttg2-6* (C), *brk1-1* (E) and *ttg2-6 brk1-1* (G). The detailed configuration of the actin arrays are highlighted in A’, C’, E’, and G’, respectively, bars=20 μm. B, D, F, and H, Representative actin arrangements in the branches of mature trichomes in WT (B), *ttg2-6* (D), *brk1-1* (F) and *ttg2-6 brk1-1* (H). The details are exhibited in B’, D’, F’, and H’, respectively, bars=20 μm. I, Quantification of anisotropy of F-actin arrays in mature trichomes in WT, *ttg2-6*, *brk1-1*, and *ttg2-6 brk1-1*. The similar parameters and methods are employed as described in Figure 4C.

To further explore the potential interaction between *TTG2* and *BRK1* during trichome development, we examined the expression level of *BRK1* in *ttg2* mutants. As shown in Figure 8A, the transcripts of *BRK1* were slightly decreased in the absence of *TTG2*. To further confirm the regulation of *BRK1* expression by *TTG2*, we generated *pBRK1:GUS* transgenic plants and introduced the transgene into *ttg2* mutant by crossing. Consistent with the quantitative RT-PCR data, the intensity of the GUS signals were much weaker in *ttg2* compared to WT (Figure 8B). Moreover, we observed that the GUS signals were mainly located in trichome cells in *pBRK1:GUS* plants (Figure 8B), indicating a high expression of *BRK1* in trichome cells. We also assessed whether transiently increasing *TTG2* transcript level could stimulate *BRK1* expression through a protoplast effector/reporter assays. The effector vector *p35S:TTG2-GR* expressed a fusion protein with the glucocorticoid receptor (GR) and TTG2 and the reporter vector was composed of two independent expression cassettes (Figure 8C). When *p35S:TTG2-GR* was co-transfected with *pBRK1:GFP- pUBQ10:mCherry*, the GFP fluorescence signals were markedly increased in DEX treated protoplasts compared with mock treatment (Figure 8, D and E), demonstrating that elevated *TTG2* could promote *BRK1* promoter activity.

**Figure 8.**
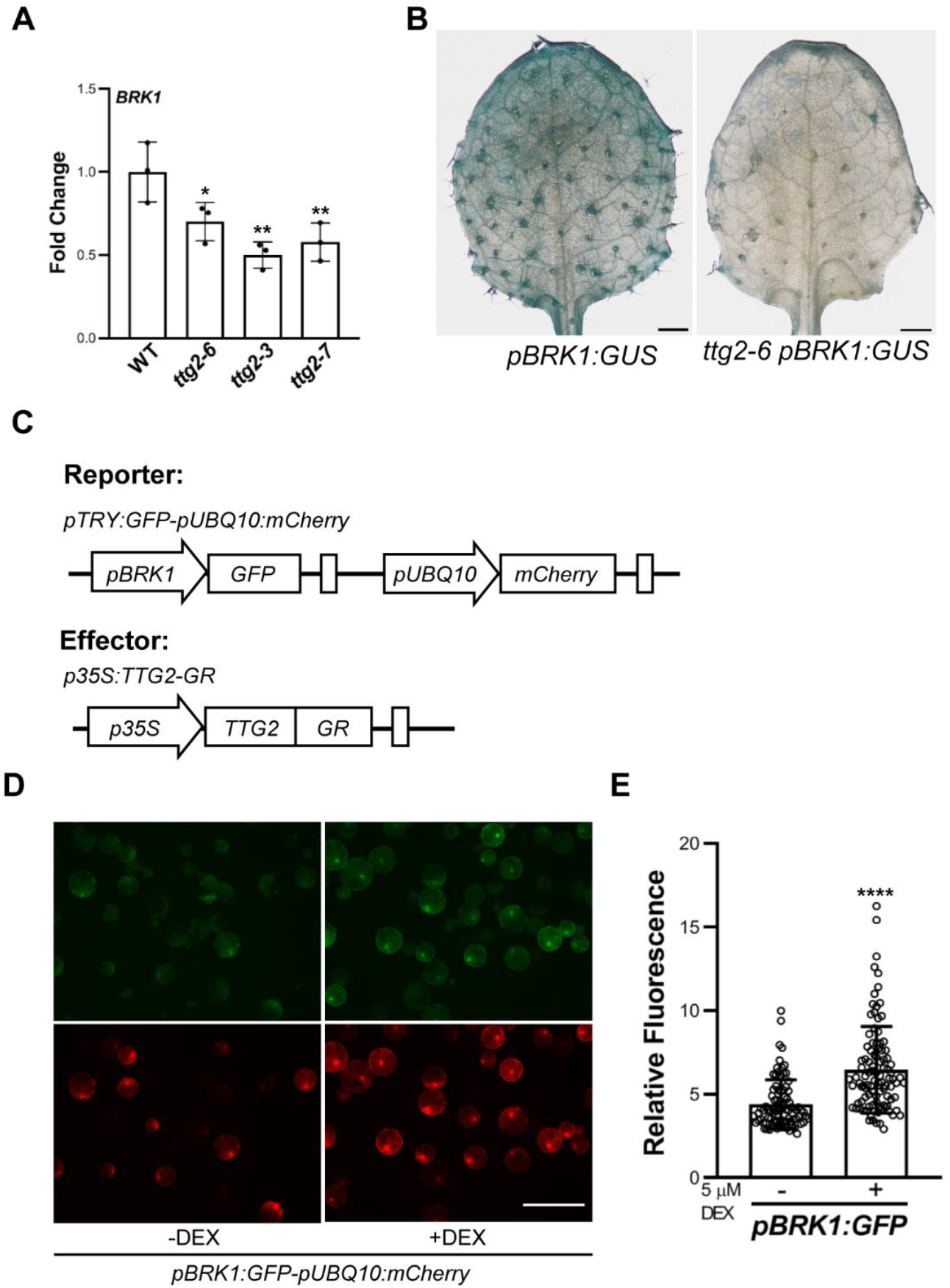
*TTG2* modulates *BRK1* expression. A, RT-qPCR analysis of the relative expression levels of *BRK1* in *ttg2* mutants. Data are shown as means ± SD of three biological replicates. **P ≤ 0.01 by the Student’s t test. B, The expression patterns of *pBRK1:GUS* in the WT and *ttg2-6*. *BRK1* expression was indicated by *GUS* expression from the fusion of the *GUS* reporter gene to the 5’ regulatory region of *BRK1*, bars=500 μm. C-E, Protoplast-based effector/reporter assays. C, Diagrams of the reporter and effector vectors. D, Representative GFP fluorescence signals in the protoplast cells when effector vector *p35S:TTG2-GR* was co-expressed with reporter vector *pBRK1:GFP-pUBQ1:mCherry* in WT mesophyll protoplasts and treated with or without DEX. GFP fluorescence was monitored using fluorescence microscopy. E, Measurements of GFP relative fluorescence intensity in protoplasts. Data are shown as means ±SD. ****P ≤ 0.0001 by the Student’s t test. Experiments were independently conducted 3 times with similar results.

Due to the positive regulation of *TTG2* on *BRK1* expression, we hypothesized that overexpression of *BRK1* may rescue the trichome defects in *ttg2* mutant. To this end, we expressed fusion protein BRK1-GFP under the control of its native promoter in *ttg2-6* mutant to generate *ttg2-6 pBRK1:BRK1-GFP* transgenic plants. The *pBRK1:BRK1-GFP* construct could rescue the developmental impairments of *brk1-1* (Supplemental Figures S8), suggesting that *BRK1-GFP* is functional *in planta*. However, the *ttg2-6 pBRK1:BRK1-GFP* plants produced unexpected trichome phenotypes. Although the trichome numbers were decreased, the overall trichome morphologies on the first pair of true leaves were similar to WT (Supplemental Figures S9, A and B). However, aberrant trichomes similar to those in *ttg2- 6* were observed on the subsequent leaves in *ttg2-6 pBRK1:BRK1-GFP* plants (Supplemental Figures S9C). These unexpected phenotypes might be due to the complicated effects of *BRK1* on trichome development, because it has been reported that overexpression of *BRK1* also caused distorted trichomes (Djakovic et al., 2006)

### TTG2 directly binds to the ***BRK1*** promoter

The genetic and molecular interactions between *TTG2* and *BRK1* prompted us to investigate whether *BRK1* functions is a direct transcriptional target of *TTG2*. Since *TTG2* has been reported to bind to its target promoters by interacting with a conserved motif (T)(T)TGAC(C/T) termed W-boxes (Pesch et al., 2014; Figure 9A), we therefore scanned the promoter region of *BRK1* for these consensus sequences. Three potential W-boxes were discovered located at - 1078‒ -1063 bp, -496‒ -487 bp, and -150‒ -141 bp (Figure 9A), and designated as W1, W2, and W3 respectively.

**Figure 9.**
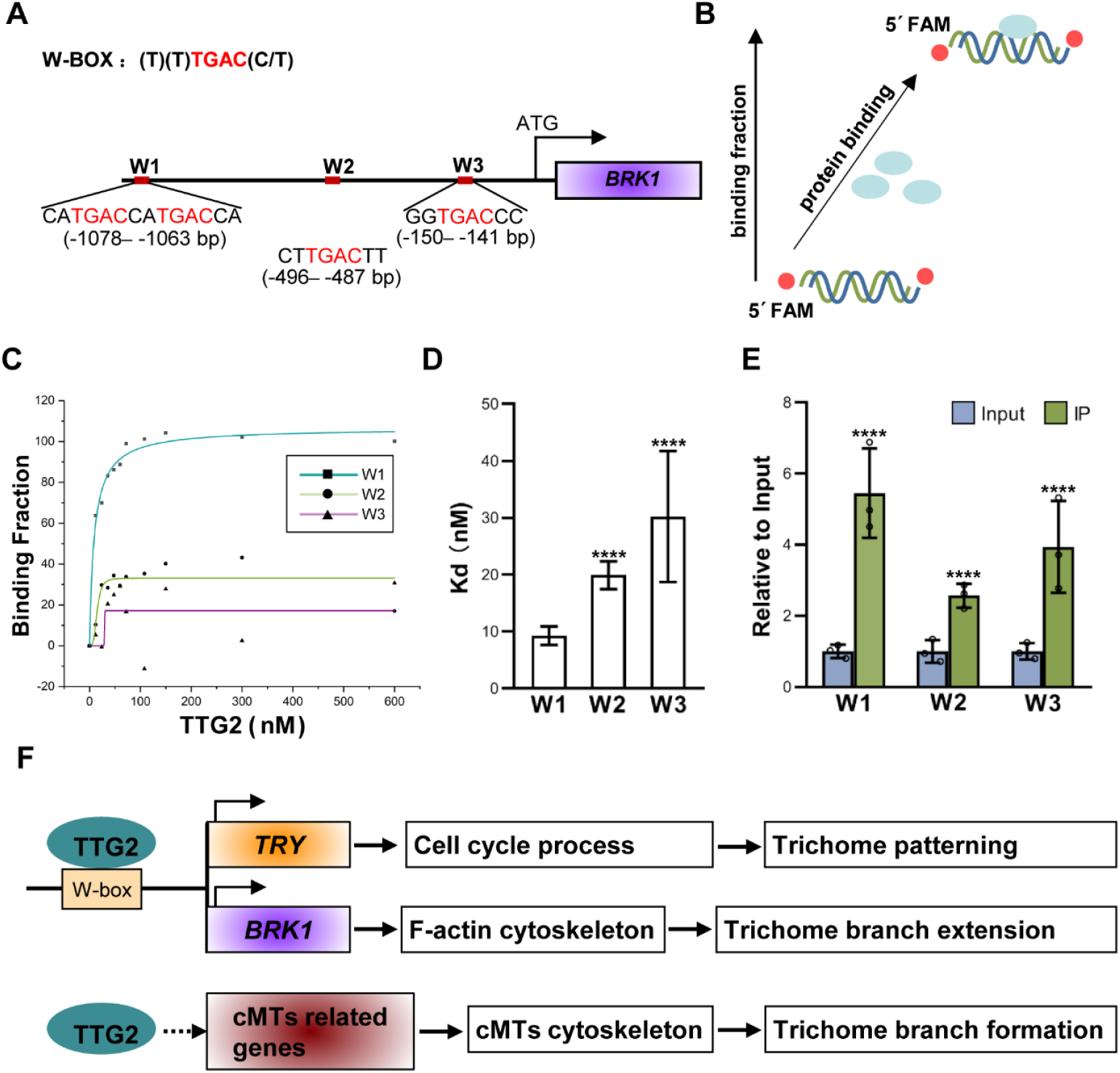
Binding of TTG2 to the *BRK1* promoter. A, The consensus sequence of the W-box, and potential W-box positions in the *BRK1* promoter. B, Schematic illustration of the FA experiment. The fluorescein FAM labeled oligonucleotide alone gives a low steady-state anisotropy value. Upon proteins (blue ovals) binding to the DNA molecule, the steady-state fluorescence anisotropy increases. The binding fraction was calculated by subtracting the steady-state fluorescence anisotropy from that when protein was added. Therefore, the direct binding of the protein and the target DNA molecular leads to an increase of binding fraction. C and D, Measurements of binding fraction (C) and the Kd (D) values in the FA experiment. The steady-state fluorescence anisotropy were determined before and after the addition of different concentrations of His-TTG2, respectively. The binding plots were simulated by using the Origin software. The Kd value were obtained through a Hill fit of the fluorescence anisotropy data. E, ChIP-qPCR analysis of the binding activities of TTG2 to W1-W3 fragments on *BRK1* promoter. The enrichment values are displayed as the ration between the input and the immunoprecipitated DNAs. The *PP2A* fragment was used as the non-binding control in qPCRs. Data were means ±SD for three biological replicates. ****P ≤ 0.0001 by the Student’s t test. F, A proposed working model for *TTG2* in controlling trichome development.

To test the direct binding of TTG2 to these potential W-boxes in the *BRK1* promoter, we first adopted an *in vitro* fluorescence anisotropy (FA) assay which is a rapid, sensitive, and real-time assay to monitor the binding between proteins and DNA fragments (Xi and Deprez, 2010). Within a given solution, the extent of depolarization depends on the rate of tumbling of the fluorescently labeled DNA molecule, which decreases with increasing size. Thus, when protein binds to a DNA molecule, accompanied by the increasing of the molecular size, the FA will decrease compared to that of the free DNA molecule (Figure 9B). According to these principles, three sequences covering each potential W-box in the *BRK1* promoter were synthesized and modified with fluorescence group FAM, respectively. In addition, the His- TTG2 recombinant proteins were expressed and purified. After measuring the value of the fluorescence anisotropy of solutions with the FAM modified DNA molecules alone and the solutions with both FAM modified DNA molecules and His-TTG2 proteins, respectively, we found a remarkable alteration of FA in the combination that contained the W1 sequence and His-TTG2 protein. Compared to the two other sequences, His-TTG2 could bind to W1 sequence in a low concentration (Figure 9C), with a lower Kd of 9.26 (Figure 9D).

Furthermore, to test if TTG2 could bind to the promoter of *BRK1 in planta*, we carried out chromatin immunoprecipitation followed by quantitative PCR (ChIP-qPCR) analyses. The above ground tissues of 2-week-old *pTTG2:TTG2-GFP* plants were harvested and ChIP was carried out with GFP-Trap agarose beads. The *pTTG2:TTG2-GFP* construct is functional since both *ttg2-6 pTTG2:TTG2-GFP* and *ttg2-7 pTTG2:TTG2-GFP* plants exhibited WT-like trichomes (Supplemental Figure S10). Consistent with the results from *in vitro* experiments, we found the W1 fragments were significantly enriched in the immunoprecipitated DNA compared with other two fragments (Figure 9E), indicating a direct binding of TTG2 to the W1 fragment in *BRK1* promoter.

Taken together, we propose a model in which *TTG2* is required at all stages during trichome development. It influences the early stage of trichome specification through its control of *TRY*. During the trichome cell morphogenesis, *TTG2* regulates proper branching by dictating cMTs configurations. During the later stages when the established pattern is elaborated, *TTG2* maintains the extension growth of the trichome stalks and branches via direct association with the actin cytoskeleton (Figure 9F).

## Discussion

### The unique trichome morphologies in ***ttg2-6***

*TTG2* encodes a zinc finger-like transcription factor of the plant-specific WRKY gene family (Johnson et al., 2002). In the first report, a major effect of *ttg2* on trichome development is to reduce trichome numbers and disrupt trichome spacing. Meanwhile, for few appeared trichomes, whose branches were reduced or eliminated (Johnson et al., 2002). Thus, *TTG2* is considered as a pivotal trichome patterning component. However, phenotype characterization of *ttg2-6* mutant in this study implying *TTG2* may be required at all stages during trichome development. One of the conspicuous phenotypes in *ttg2-6* is the presence of varied branched trichomes (Figure 1, H and I). Such a trichome phenotype is really interesting, because to date, almost all of the trichome branching defective mutants reported display consistently branching phenotypes, with either less branched or over branched trichome (Tominaga-wada et al., 2011; Han et al., 2022). Moreover, the trichome phenotypes of *ttg2-6* also seems inconsistent with other *ttg2* alleles both whatever in L*er* or in Col background (Johnson et al., 2002; Ishida et al., 2007; Figure 3). One of the possible explanations is that *ttg2-6* represents a weak mutant allele of *TTG2*. First, the overall trichome branch numbers are reduced in *ttg2-6* (Figure 1, H and I), which is inconsistent with other *ttg2* alleles (Johnson et al., 2002; Ishida et al., 2007; Figure 3). Second, like that in *ttg2-3*, the expression pattern of *CPC*, *GL2*, and *TRY* were also altered in *ttg2-6* mutant (Supplemental Figure S11). Third, in contrast to other *ttg2* mutants, the mutation occurs at the very C-terminal in *ttg2-6* (Figure 2, A and B), which may allow partial functions of *TTG2* to be preserved. Meanwhile, given mutation in *ttg2-6* resulted in the loss of one of the two conserved Cys residues in the second putative zinc finger motif (Johnson et al., 2002; Figure 2A), the Cys residue therefore might be critical for *TTG2* in regulating trichome branching. In addition, although it has been reported that some *ttg2-1* trichomes show a range of distorted growth patterns (Johnson et al., 2002), which is not as remarkable as that in *ttg2-6* (Figure 1, B, E, and G).

Collectively, the isolation of the *ttg2-6* mutant provides a distinct genetic material to further unravel the molecular function of *TTG2* during trichome development.

### ***TTG2*** integrates microtubule cytoskeletal and cell cycle signals to control trichome branching

Accumulated information have demonstrated cell cycle control and microtubule organization are two essential pathways required for trichome branch initiation (Mathur and Chua, 2000; Schnittger and Hülskamp, 2002; Ishida et al., 2008; Tominaga-Wada et al., 2011). Generally, the nuclear DNA content of mature trichomes will reach to 32C (C equals haploid DNA content per nucleus), accompanied by the formation of three or four branches (Schnittger and Hülskamp, 2002). Mutations in genes that increase or decrease the endoreduplication levels elevate or reduce the nuclei size and the number of branches respectively (Tominaga-Wada et al., 2011; Han et al., 2022). Meanwhile, during trichome morphogenesis, the arrangement of cMTs changes dramatically at branching points (Mathur and Chua, 2000). Mutation in genes involved in the cMTs assemble or dynamics often lead to aberrant trichome branching (Tominaga-Wada et al., 2011; Li et al., 2019). Given the production of remarkable over branched trichomes in *ttg2-6* (Figure 1, B, and E-I), we examined the trichome nuclear size in *ttg2-6* and revealed an apparent increased nuclei size and nuclear DNA content (Supplemental Figure S12, A-C). We also verified global DNA ploidy levels in *ttg2-6* plants with flow cytometry. Compared to that in WT, a marked shift of cells to higher ploidy levels was observed in *ttg2-6*, with a significantly higher frequency of 16C cells and the presence of 32C cells which were not seen in the WT (Supplemental Figure S12D). In consistent, the *ttg2- 6* plants displayed enlarged cotyledons (Supplemental Figure S12, E, G) and cotyledon pavement cell sizes compared with those of the WT (Supplemental Figure S12, F, H). These findings suggest that *TTG2* may be involved in the endoreduplication regulation during trichome morphogenesis. Conincidentally, *TTG2* was reported to directly activate the expression of *TRY* that regulates trichome morphologies in a cell cycle-dependent manner (Szymanski and Marks, 1998). Therefore, *TRY* may mediate the regulation of *TTG2* on cell cycle control.

Meanwhile, the cMTs arrangements are obviously disrupted (Figure 4), establishing the regulation of *TTG2* on the microtubule cytoskeleton. To elaborate the underlying mechanisms of the regulations of *TTG2* on cMTs, we crossed *ttg2-6* with those trichome branching mutants associated with cMTs assembly or dynamics reported before to implement genetic interaction analysis. Additive effects were observed by examining the trichome phenotypes of different combinations (Supplemental Figures 5 and 6), suggesting *TTG2* may act in cooperating with these genetic factors to regulate microtubule cytoskeleton during trichome development.

Therefore, other strategies liking RNA-seq or ChIP-seq could be employed to screen the potential *TTG2* targets associated with cMTs cytoskeletal pathway.

### ***TTG2*** regulates trichome branching expansion through directly interacting with actin cytoskeletal component

In addition to trichome branching defects, the branching expansion is also impaired due to *TTG2* mutation. A proportion of trichomes in *ttg2-6* display striking distorted phenotypes (Figure 1, E and G), resembling to that of the SCAR/WAVE and ARP2/3 mutants (Hülskamp et al., 1994; Schwab et al., 2003). This phenotypic association implies the function of *TTG2* on actin cytoskeletal organization during trichome branching expansion (Schwab et al., 2003; Hülskamp, 2004). As respected, we observed the F-actin organization is dramatically disrupted in *ttg2-6* trichomes (Figure 5).

*BRK1* encodes a member of SCAR/WAVE complex and participates in ARP2/3 activation (Gallagher and Smith, 2000; Frank and Smith, 2002; Djakovic et al., 2006; Le et al., 2006). Mutation of *BRK1* results in distorted trichome morphology and aberrant F-actin cytoskeleton organization (Gallagher and Smith, 2000; Frank and Smith, 2002; Djakovic et al., 2006).

Genetic analyses indicate that *brk1* is epistasis to *ttg2-6* with regard to branching expansion (Figure 6B). Moreover, the expression of *BRK1* is tightly regulated by *TTG2* (Figure 8), indicating *BRK1* may serve as a potential target of *TTG2*. Our further biochemical and molecular experiments showed *TTG2* could directly bind to a W-box in the *BRK1* promoter (Figure 9). These findings suggest the effect of *TTG2* on actin cytoskeleton organization may be achieved through modulating the expression of *BRK1*. In addition, we found the transcript accumulation of *BRK1* is apparently inhibited in *gl1*, *ttg1*, and *gl3* mutants, and the trichome phenotypes of *gl1 brk1-1*, *ttg1 brk1-1* and *gl3 brk1-1* double mutants resemble the *brk1-1* single mutant respectively (Supplemental Figure 13). These preliminary results suggest *BRK1* may act downstream of the MYB-bHLH-WD-40 transcription activation complex.

## Conclusions

Taking advantages of the unique trichome phenotypes in *ttg2-6*, we uncover the new functions of *TTG2* and suggest that *TTG2* is required at all stages of during trichome development. In addition, our studies directly link *TTG2* and cytoskeletal component, providing new insight into cellular signaling events downstream of the transcriptional factors associated with trichome development in Arabidopsis.

## Material and methods

### Plant materials and growth conditions

Plant materials used in this study are all in the *Columbia-0* (*Col-0*) background except for the map-based cloning. The wide type (WT) refers to *Col-0* plants. T-DNA insertion lines for *TTG2* (*Salk_148838*, *ttg2-3*; *Salk_206852*, *ttg2-7*) (Ishida et al., 2007), *BRK1* (*CS86554*, *brk1-1*; *CS93199*, *brk1-2*) (Djakovic et al., 2006), *ZWI* (*Salk_017886*, *zwi-101*) (Chen et al., 2016), *AN* (*Salk_026489*) (Gachomo et al., 2013), *KTN1*(*Sail_343_D12*) (Chen et al., 2014), *CROOKED* (*Salk_123936*) (Bellinvia et al., 2022), *WRM* (*Salk_003448*) (Le et al., 2003), *DIS2* (*Salk_201281*), *IBT* (*Salk_039449*) (Uhrig et al., 2007), *GRL* (*Salk_135634*) (El-Assal et al., 2004), *GL1* (*SALK_133117*), *TTG1* (*CS67772*, *ttg1-13*), *gl3-1* (Koornneef et al., 1982), and a marker line for microtubule array *GFP-TUB6* (*CS6550*) (Liang et al., 2019) were obtained from the Arabidopsis Biological Resource Center (ABRC). The *ttg1-13* and *gl3-1* lines were crossed three times to *Col-0* respectively before used for construction double mutants. Double mutants were constructed by crossing. The T-DNA insertion sites and homozygous plants were confirmed by genomic PCR and DNA sequencing. The marker line for actin filaments *ABD2-GFP* was kindly provide by Prof. Zhaosheng Kong (Chinese Academy of Sciences). Primers used for genotyping were listed in Supplemental Table 1.

For plant culture, seeds were stratified at 4°C for 2 d and then sowed on commercial soil mix (Pindstrup) for germinating and growth in growth room at 22±1°C under continuous illumination (∼100 μmol m^−2^ s^−1^). For protoplast preparation, seeds were sowed and grown on Jiffy-7-Peat Pellets (Jiffy Group) in a growth chamber (Conviron A1000) with day/night cycle (12h/12h) at 22±1°C, and fully expanded rosette leaves of 30 d old plants were used.

### Trichome phenotype characterization

Trichome phenotypes were examined as described by Liang et al., (2019). Briefly, plants in different genotype backgrounds were grown in soil for three weeks and trichome morphology of the 3^rd^ and 4^th^ rosette leaves were examined under stereoscope (SZ61, Olympus, Japan). The branch numbers, clusters, and the trichome density were counted and calculated.

Meanwhile, representative trichomes on the 5^th^ or 6^th^ rosette leaves were imaged by using a tabletop scanning electron microscopy (SEM TM3030, Hitachi, Japan). For each genotype, at least 15 individuals were used for quantitative analysis, and all experiments were repeated for three times.

For all quantitative analyses, student’s t-test was used to assess the statistical significance.

### Mutant screening and map-based cloning

The *abt4-1* (*ttg2-6*) mutant was isolated from an ethyl methanesulfonate (EMS)-mutagenesis mutant population in *gl2-3* mutant background (*Salk_039825,* Wang et al., 2010) we established before. The *abt4-1 gl2-3* double mutants was initially isolated with transparent and distorted trichome. The *abt4-1* single mutant was obtained by backcrossing the *abt4-1 gl2-3* mutant with *Col-0* plants, and two additional rounds of backcross were carried out before phenotypic characterization.

The mutation site in *abt4-1* was determined via map-based cloning according to Lukowitz et al. (2000). In brief, *abt4-1* was crossed with Arabidopsis Landsberg *erecta* (L*er*) ecotype to generate an F2 mapping population. Then, bulked segregant analysis (BSA) with DNA pool mixed from 95 individuals from the mapping population and 25 pairs of molecular markers distributed on the five chromosomes was carried out to determine the chromosomal region that is linked to *ABT4*. After that, additional molecular markers were used to fine map and narrow down the linked region. Lastly, candidate genes in the linked region were sequenced to finally confirm the mutation site. Sequence information of the molecular markers used were listed in the Supplemental Table S1.

### Generation of transgenic plants

To conduct complementation experiments, the genomic fragment comprising its 1,086 bp endogenous promoter and the full-length coding region of *TTG2* were amplified and cloned into the *pCambia1300* vector to generate *pTTG2:TTG2* plasmid. To evaluate the effects of *BRK1* on trichome development in *ttg2* mutant background, the *pBRK1:BRK1-GFP* construct was generated. The GFP gene was amplified and cloned into the *pCambia1300* vector to generate *p35S:GFP* plasmid firstly, then the genomic fragment of the *BRK1* including the 1,515 bp promoter sequences was amplified and cloned into the upstream of *GFP* to generate *pBRK1:BRK1-GFP* plasmid. *pTTG2:TTG2-GFP* construct was generated by using the similar procedures to perform the ChIP experiment. *pBRK1:GUS*, *pCPC:GUS*, *pGL2:GUS*, *pTRY:GUS*, plasmid was obtained by introducing the promoter sequence of *BRK1*, *CPC*, *GL2*, and *TRY* into the upstream of *GUS* gene in *pCB308* vector respectively. All the primers used were listed in Supplemental Table S1.

The confirmed constructs were transformed into the indicated genetic background via Agrobacterium mediated floral-dip method (Clough and Bent, 1998). Transgenic plants were obtained through antibiotic-resistant, genomic PCR, and trichome phenotype examination, respectively. The T3 plants were used for phenotype characterization and other experiments. At least 20 independent transgenic lines were obtained for each transformation, and two representatives were used for detailed analyses.

### RNA isolation and quantitative RT-PCR

Total RNAs were extracted from the 5^th^-7^th^ rosette leaves of 2-week-old soil grown plants with TRIzol reagent (15596026, ThermoFisher Scientific). cDNAs were obtained with oligo (dT15) primer by using the Maxima H Minus cDNA Synthesis Master Mix (M1662, Thermo Fisher Scientific). Real-time PCRs were carried out with the Fast Start Essential DNA Green Master kit (06402712001, Roche) and Bio-Rad CFX96 real-time PCR system. Relative expression levels of the target genes were calculated with 2^-△Ct^. Expressions of *ACTIN2* (*ACT2*) was used as the internal controls. Primers for RT-qPCRs were listed in Supplemental Table S1.

### Cortical microtubules (cMTs) and F-actin microfilaments imaging and quantification

To achieve *in planta* cortical microtubules (cMTs) and actin filaments (F-acin) observation, marker lines *GFP-TUB6* and *ABD2-GFP* were crossed to the different mutant background respectively and the double mutants were selected by genomic PCR and GFP fluorescence examination. Trichomes on the first pair of true leaves from 10-day-old soil grown seedlings were mounted and examined under a spinning disk confocal system built on a DMi8 inverted microscope (Leica) equipped with a CSU-W1 confocal scanner unit (Yokogawa) and an iXon Ultra 888 EMCCD camera (Andor). cMTs and F-actin were observed with a HCPLAPO 63x or 100x N.A.1.30 glycerol objective (Leica) and images were taken through Z-stacking with a 0.5 um z-step.

cMTs and F-actin angles and anisotropy quantification were performed according to Yao et al. (2008). Digital images acquired were loaded in Fiji-ImageJ and angles of individual cMTs and F-actin were measured respectively. The value of microtubules angles of that parallel to the trichome branch or stalk extension directions were defined as 0°, while the value of those were perpendicular to the cell’s longitudinal axis were defined as 90°. The cMTs and F-actin anisotropy were evaluated according to Boudaoud et al. (2014), and the following convention was used: the anisotropy score 0 for no order (purely isotropic arrays) and 1 for perfectly ordered (purely anisotropic arrays).

### Protoplast effector/reporter assays

The experimental procedures were conducted according to Iwata et al. (2011). The reporter plasmid *pBRK1:GFP-pUBQ10:mCherry* and effector plasmid *p35S:TTG2-GR* were constructed, respectively. Primers used were listed in Supplemental Table S1. The mesophyll protoplast from WT were prepared and reporter and effector plasmids (20 μg each) were co-transformed into 200 μl mesophyll protoplasts according to Yoo et al.(2007). After transfection, protoplasts were treated with 5 μM dexamethasone (DEX) or equal volume of ethanol and incubated in W5 buffer (154 mM NaCl, 125 mM CaCl2, 5 mM KCl, 2 mM MES) for 12 h. The fluorescence signals were examined under a fluorescence microscope (DMI8, Leica) and mCherry fluorescence was used as the transfection control. For each transfection, more than 100 protoplasts which showed mCherry signals were used to measure GFP fluorescence signal intensity with Image J software. The experiments were repeated for three times with similar results.

### Fluorescence anisotropy (FA) experiment

To test the direct binding of TTG2 to the promoter of *BRK1*, the FA experiment were employed according to Xi and Deprez (2010). First, the recombined protein His-TTG2 was obtained by inducing expression of *pET28a-His-TTG2* in *E. coli* strain Rosetta (DE3) and purifying soluble proteins through affinity chromatography with Ni^2+^-nitrilotriacetate acid columns followed by fast protein liquid chromatography with size exclusion chromatography columns according to manufacturer’s specifications. Primers used were listed in Supplemental Table S1. Secondly, the fragments which carry the potential W-box in the *BRK1* promoter were synthesis and modified with fluorescence group FAM at the 3’ and 5’ end, respectively. Thirdly, the FAM modified DNA fragments were annealed and the steady-state fluorescence anisotropy of the double strand DNA was evaluated with a commercial fluorescence polarization system (Beacon 2000 Fluorescence Polarization, PanVera Corporation). Lastly, the His-TTG2 protein was added at the indicated concentrations, and the fluorescence anisotropy value of the solution was measured. The binding fraction is calculated by subtracting the steady-state fluorescence anisotropy from that when His-TTG2 protein was added. The dissociation constant (Kd) were calculated according to 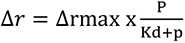, where P corresponds to protein concentration, △rmax represents the maximal fluorescence anisotropy difference value.

### Chromatin immunoprecipitation (ChIP) experiment

ChIP experiments were performed as described by An et al. (2012). In short, 3-4 g of the 5^th^- 7^th^ rosette leaves from 2-week-old soil grown *pTTG2:TTG2-GFP* plants were harvested and fixed with crosslinking buffer (10mM Tris-HCl pH 8.0, 0.4 M sucrose, 10 mM MgCl2, 5 mM β- ME, 1 mM PMSF, and 1% formaldehyde (v/v)) for 10 min under vacuum. The treated tissues were ground in liquid nitrogen and the nuclei were extracted and collected with extraction buffer (0.4 M sucrose, 10 mM Tris-HCl pH 8.0, 10 mM MgCl2, 5mM mercaptoethanol, 0.1 mM phenylmethanesulfonyl fluoride, 1x protease inhibitor) and nuclear lysis buffer (50 mM Tris-HCl pH 8.0, 10 mM EDTA, 1% (w/v) SDS, 1 mM PMSF, 1X protease inhibitor cocktail). The lysate was sonicated with Bioruptor Pico (Diagenode) to shear the chromatin, and the resulting solution was incubated with GFP-Trap agarose beads (gta-20, Chromo Tek) at 4°C overnight with gentle shaking to immunoprecipitate chromatin complexes. The immunoprecipitated DNAs were obtained from the complexes through reverse crosslinking, proteinase K (EO0491, Thermo Fisher Scientific) digestion, and purification with QIAquick PCR Purification Kit (28104, Qiagen). DNA levels in input and immunoprecipitated solutions were determined by qPCR and fold enrichments was calculated as the ration between the input and the immunoprecipitated DNAs. A PP2A fragment was used as the non-binding control in qPCRs. Primers used are listed in Supplemental Table S1. The experiments were repeated for three times with similar trends.

## Accession number

Sequence data from this article can be found in the TAIR/GenBank data base with the accession numbers showed in Supplemental Table S2.

## Supplemental data

Supplemental Table S1. Primer pairs used in this work.

Supplemental Table S2. Accession numbers for the genes cited in this work.

## Supplemental experimental procedures

**Supplemental Figure S1.** The overall plant morphology of *abt4-1*.

**Supplemental Figure S2.** Map-based cloning of the *ABT4* locus.

**Supplemental Figure S3.** Arrangements of cMTs and F-actin in *ttg2-6* cotyledons.

**Supplemental Figure S4.** Trichome phenotypes of the *ttg2-6 zwi-101* plants.

**Supplemental Figure S5.** Trichome phenotypes of the *ttg2-6 an* and *ttg2-6 ktn1-1* plants.

**Supplemental Figure S6.** Trichome phenotypes of the double mutant combined with *ttg2-6* and different *ARP2/3* mutants.

**Supplemental Figure S7.** Trichome phenotypes of the double mutant combined with *ttg2-* and different *SCAR/WAVE* mutants.

**Supplemental Figure S8.** Complementation of the *brk1-1* mutant with the *pBRK1:BRK1-GFP* construct.

**Supplemental Figure S9.** Trichome phenotypes of *ttg2-6 pBRK1:BRK1-GFP* plants.

**Supplemental Figure S10.** Complementation of the *ttg2* mutants with the *pTTG2:TTG2-GFP* construct.

**Supplemental Figure S11.** Expression patterns of *CPC*, *GL2* and *TRY* in *ttg2-6*.

**Supplemental Figure S12.** Characterization of DNA ploidy in *ttg2-6*.

**Supplemental Figure S13.** Genetic analysis of the MYB-bHLH-WD-40 transcription complex and *BRK1* in trichome morphogenesis.

## Acknowledgments

This work is supported by the National Natural Science Foundation of China (31470290, 32070198), the Chinese Universities Scientific Fund (2452020180), and the U.S. National Science Foundation (1923589 and IOS-2127485). We also thank Prof. Hou (Northwest A&F University) for technical support for FA experiment.

## Conflict of interest statement

The authors declare no conflict of interest.

## Author contributions

LA and LL conceived the project, and arranged the experiments. LL, CW, YL, LY, WY, CZ, DW performed the experiments; LA, LL and WW analyzed the data; LL and LA wrote the manuscript; HZ, FY, and JS revised the manuscript.

